# Cell migration simulator-based biomarkers for glioblastoma

**DOI:** 10.1101/2023.02.24.529880

**Authors:** Jay Hou, Mariah McMahon, Jann N. Sarkaria, Clark C. Chen, David J. Odde

## Abstract

Glioblastoma is the most aggressive malignant brain tumor with poor survival due to its invasive nature driven by cell migration, with unclear linkage to transcriptomic information. Here, we applied a physics-based motor-clutch model, a cell migration simulator (CMS), to parameterize the migration of glioblastoma cells and define physical biomarkers on a patient-by-patient basis. We reduced the 11-dimensional parameter space of the CMS into 3D to identify three principal physical parameters that govern cell migration: motor number – describing myosin II activity, clutch number – describing adhesion level, and F-actin polymerization rate. Experimentally, we found that glioblastoma patient-derived (xenograft) (PD(X)) cell lines across mesenchymal (MES), proneural (PN), classical (CL) subtypes and two institutions (N=13 patients) had optimal motility and traction force on stiffnesses around 9.3kPa, with otherwise heterogeneous and uncorrelated motility, traction, and F-actin flow. By contrast, with the CMS parameterization, we found glioblastoma cells consistently had balanced motor/clutch ratios to enable effective migration, and that MES cells had higher actin polymerization rates resulting in higher motility. The CMS also predicted differential sensitivity to cytoskeletal drugs between patients. Finally, we identified 11 genes that correlated with the physical parameters, suggesting that transcriptomic data alone could potentially predict the mechanics and speed of glioblastoma cell migration. Overall, we describe a general physics-based framework for parameterizing individual glioblastoma patients and connecting to clinical transcriptomic data, that can potentially be used to develop patient-specific anti-migratory therapeutic strategies generally.

**Significance Statement:** Successful precision medicine requires biomarkers to define patient states and identify personalized treatments. While biomarkers are generally based on expression levels of protein and/or RNA, we ultimately seek to alter fundamental cell behaviors such as cell migration, which drives tumor invasion and metastasis. Our study defines a new approach for using biophysics-based models to define mechanical biomarkers that can be used to identify patient-specific anti-migratory therapeutic strategies.

## Introduction

Glioblastoma is the most common malignant brain tumor with median survival of only 15 months and less than 5% five-year survival rate [1,2]. Complete surgical resection is difficult because the tumor is highly invasive, and tumor cell infiltration toward the surrounding brain tissue drives disease progression and recurrence [3,4]. Glioblastoma patient-derived xenograft (PDX) and patient-derived (PD) cells (collectively referred to here as “PD(X)”) can retain the genomic profiles and histopathology of parental tumors [5–8], and have been used extensively to study glioblastoma cell migration and invasion [9–12]. However, in these studies, the connection of experimental observations to fundamental glioma cell migration mechanics as a function of the transcriptomically-defined glioblastoma subtypes of proneural (PN), classical (CL), and mesenchymal (MES) [13–15] was still unclear. In addition, it is not always feasible clinically to conduct *in vitro* migration assays on patient cells, and different harvesting and culturing methods may significantly alter the migration behavior [16]. Therefore, to effectively target cancer cell migration, it is critical to understand the fundamental mechanics of glioblastoma cell migration and its link to transcriptomic information to predict tumor cell invasion based on patient-specific omic analysis.

In the classic cell migration cycle, the first step is the extension of a cell protrusion at the leading edge driven by actin polymerization into self-assembled actin filaments (F-actin). F-actin undergoes retrograde flow driven by myosin II (motor)-mediated contraction, leading to protrusion retraction. At the same time, cell adhesion molecule binding to the extracellular environment, and subsequent stretching of the actin-adhesion adaptor proteins, constitute a molecular “clutch” that resists myosin forces and biases the protrusion toward net extension. The adhesion proteins can form focal adhesions that allow the cell to transmit traction forces onto compliant substrates. This system is known as the motor-clutch mechanism and is widely-used to describe cell migration [17–19]. Stochastic simulations of the motor-clutch model [20] have been developed and applied to predict the cell traction force, cell morphology, and F-actin flow on various substrates [21–26]. Beyond single protrusions, the cell nucleates multiple protrusions via F-actin polymerization, each of which can be modeled as a motor-clutch system, with traction forces balancing across the different protrusions. Stochastic perturbations to the force balance due to adhesion bond rupture enable larger scale cell movements and can define the front and the rear of the cell [27,28]. By imposing a force balance between the protrusions, Bangasser et al., 2017 [26] successfully developed a whole-cell motor-clutch model, the cell migration simulator (CMS, Figure 1a), to simulate the optimal cell migration of U251 glioma cells on polyacrylamide (PA) gels of different stiffnesses, and subsequent work predicted the optimal cell motility with different focal adhesion sizes and distributions [29], simulated the higher cancer cell motility on viscoelastic gels with faster stress relaxation [30], predicted the negative durotaxis for U251 glioma cells when the cells migrated toward the softer region of the PA gel with stiffness gradients [31], and simulated the cyclic cell migration speed within 1D channels for melanoma cells [32]. By changing the parameter values, the CMS can capture the different migration features in different cell types and under drug treatments, which may link to cell transcriptomes [26]. Therefore, the CMS provides a consistent mechanical framework which can potentially be used to interpret and synthesize cell migration and force measurements across glioblastoma PD(X) systems and subtypes in order to predict cell migration based on glioblastoma cell transcriptomes.

**Figure 1.**
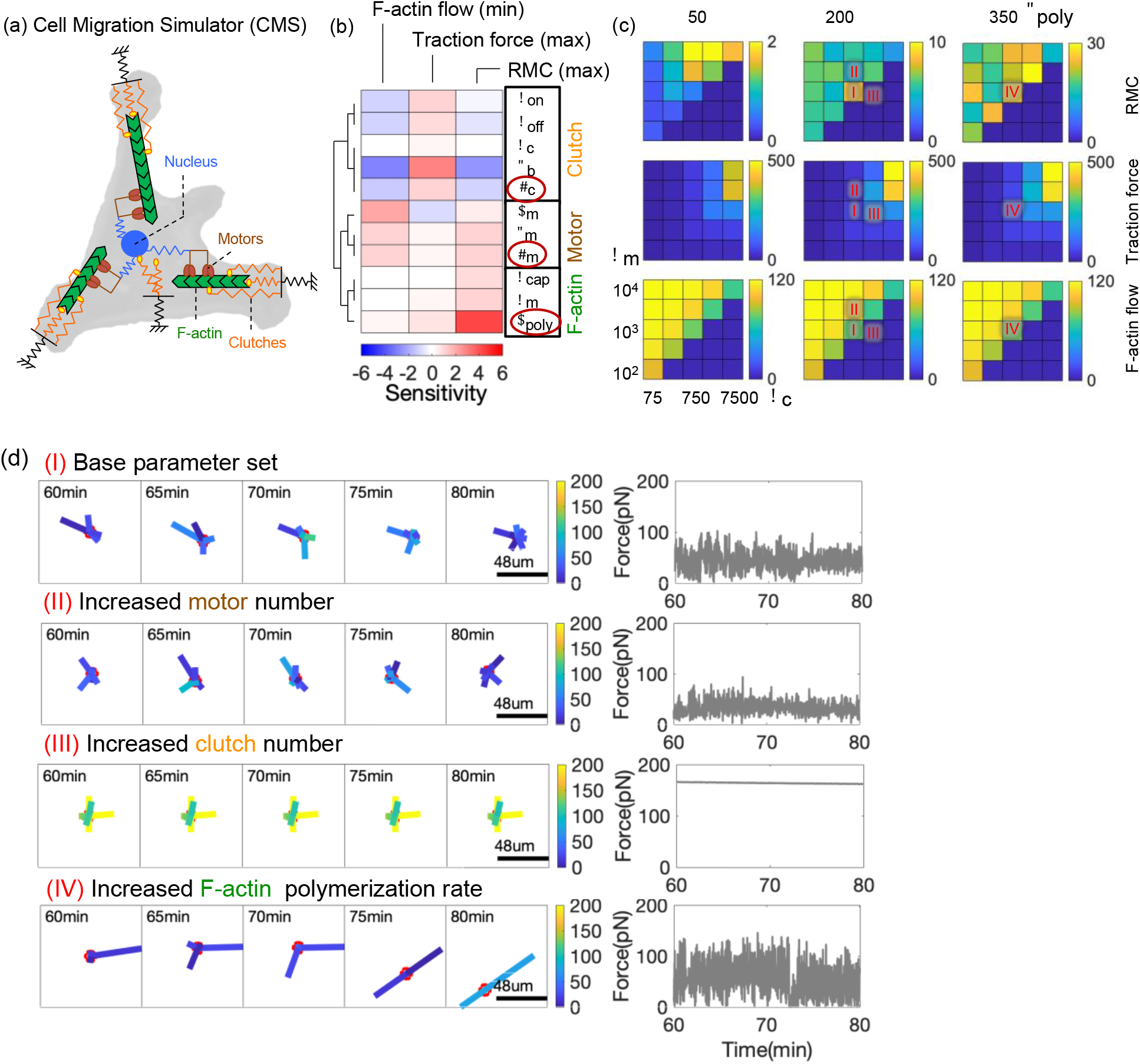
Three CMS physical parameters dictate cell migration and traction force. (a) CMS schematic. (b) Parameter sensitivities of the CMS were analyzed to predict maximum cell migration speed (RMC), minimum F-actin flow, and maximum traction force across substrate stiffnesses. With hierarchical clustering, three parameter groups were identified: clutch group (*n*_c_, *F*_b_, *k*_on_, *k*_off_), motor group (*n*_m_, *F*_m_, *v*_m_) and actin group (*v*_poly_, *k*_mod_, *k*_cap_). The three parameters (*n*_m_, *n*_c_, *v*_poly_) were chosen as fundamental physical expressions of the CMS. (c) Maximum RMC, maximum traction, minimum F-actin flow across substrate stiffnesses as a function of the three biophysical expressions (*n*_m_, *n*_c_, *v*_poly_) predicted by the CMS were plotted. Condition I, II, III, IV represent distinct cell migratory behaviors with different (*n*_m_, *n*_c_, *v*_poly_). (d) Condition I represents a typical migrating cell with base parameter values. In Condition II, higher motor number resulted in lower traction, shorter protrusion length, and slower cell migration. In Condition III, higher clutch number resulted in a near-maximal constant traction, limited dynamic protrusions, and poor cell migration. In Condition IV, higher actin polymerization rate resulted in more dynamic protrusions, longer protrusion length, highly fluctuating traction, and faster cell migration.

In this study, we explored the ability of the CMS to serve as a physics-based framework for glioblastoma subtypes and PD(X) systems. We used the CMS parameters representing myosin II motors, adhesion protein “clutches,” and F-actin polymerization to predict cell migration generally, and then mechanically parameterized glioblastoma cells obtained from a cohort of 11 glioblastoma patients across all three subtypes and two different culture procedures. Using single cell migration and force generation data obtained on compliant 2D surfaces, we found distinct parameter sets for glioblastoma patients across subtypes and culture conditions. In addition, the CMS predicted differential cell migration sensitivities to cytoskeletal drugs between subtypes. Finally, we established correlative links between the CMS parameter values and patient transcriptomes. Our results suggest it is feasible to estimate cell migration speeds using mRNA expression, similar to how migration speed can be estimated via machine learning-based detection of features in clinical MRI images [33]. Overall, we describe a consistent physics-based framework for parameterizing individual glioblastoma patients, connected to clinical transcriptomic data, that can potentially be used to develop subtype and patient-specific anti-migratory therapeutic strategies for glioblastoma and potentially other high-grade malignancies.

## Results

### Three CMS physical parameters dictate cell migration and traction force

The CMS, shown schematically in Figure 1a, can reproduce the experimentally observed optimal cell migration on compliant PA gels in the range of *in vivo* tissue stiffnesses (~1-200 kPa) (Figure 1 in Bangasser et al., 2017 [26]). To understand the relationship between the CMS parameters and its predictions, we computed the optimal cell motility (random motility coefficient, RMC), traction force, and F-actin retrograde flow in the range of substrate stiffnesses across a wide range of parameter values (Figure S1). Similar to the analysis in Bangasser et al., 2013 [21], the sensitivities of the fold changes in the CMS predictions to the fold changes in parameter values, referred to as the parameter sensitivities, were plotted in Figure 1b. Results showed that clutch-related parameters (*n*_c_, *F*_b_, *k*_on_, *k*_off_, *k*_c_, Table 1) increased the cell traction force and reduced the F-actin flow, motor-related parameters (*n*_m_, *F*_m_, *v*_m_, Table 1) increased the F-actin flow and motility, and actin-related parameters (*k*_mod_, *v*_poly_, *k*_cap_, Table 1) increased the motility (Figure 1b). We applied unsupervised hierarchical clustering to the parameter sensitivities, and found that the motor, clutch, and actin-related parameters naturally clustered into motor, clutch, and actin groups, respectively (Figure 1b). Therefore, the CMS parameters can be categorized broadly into three groups, and each has its own unique influence on predicted cell migration, which allows us to reduce the 11-dimensional parameter space to 3 fundamental dimensions of motor, clutch, and actin parameters.

**Table 1.** Parameters for the cellular level CMS.

| Parameter | Symbol | Value | Ref. |
| --- | --- | --- | --- |
| Total number of myosin motors | $n_m$ | 1,000 | Adjusted |
| Total number of clutches | $n_c$ | 750 | Adjusted |
| Maximum total actin length | $A_{\text{tot}}$ | 100 $\mu\text{m}$ | 26 |
| Maximum actin polymerization rate | $v_{\text{poly}}$ | 200 nm/s | Adjusted |
| Maximum module nucleation rate | $k_{\text{mod}}$ | 1 $\text{s}^{-1}$ | 26 |
| Module capping rate | $k_{\text{cap}}$ | 0.001 $\text{s}^{-1}$ | 26 |
| Initial module length | $l_{\text{in}}$ | 5 $\mu\text{m}$ | 26 |
| Minimum module length | $l_{\text{min}}$ | 0.1 $\mu\text{m}$ | 26 |
| Cell spring constant | $k_{\text{cell}}$ | 10,000 pN/nm | 26 |
| Number of cell body clutches | $n_{\text{c,cell}}$ | 10 | 26 |
| Substrate spring constant | $k_s$ | 0.3–300 pN/nm | Adjusted |
| Maximum number of module motors | $n_m$ | 1,000 | 26 |
| Myosin motor stall force | $F_m$ | 2 pN | 26 |
| Unloaded actin flow rate | $v_m$ | 120 nm/s | 26 |
| Maximum number of module clutches | $n_c$ | 750 | 26 |
| Clutch on-rate | $k_{\text{on}}$ | 1 $\text{s}^{-1}$ | 26 |
| Unloaded clutch off-rate | $k_{\text{off}}$ | 0.1 $\text{s}^{-1}$ | 26 |
| Clutch spring constant | $k_c$ | 0.8 pN/nm | 26 |
| Characteristic clutch rupture force | $F_b$ | 2 pN | 26 |

Because of the natural clustering into three distinct groups, we chose one parameter from each clustered group as the fundamental physical parameters: motor number (*n*_m_) representing myosin II motor activity, clutch number (*n*_c_) representing functional adhesion protein level, and Factin polymerization rate (*v*_poly_) representing actin filament polymerization activity. As described previously, these three parameters are the key components in the cell migration process [17–19], they exhibit different values in different cell types [27,28], and they are easily manipulated by drugs [26]. Therefore, we used these three parameters (*n*_m_, *n*_c_, *v*_poly_) as a fundamental basis set to predict glioblastoma cell migration across subtypes and culture conditions in the absence and presence of drugs.

To illustrate the cell migration governed by the three fundamental physical parameters, we plotted the optimal motility (RMC), traction force, and F-actin flow as a function of the three physical parameters (*n*_m_, *n*_c_, *v*_poly_) (Figure 1c). There are four different scenarios observed in the parameter space (Figure 1c, Condition I, II, III, IV), and the time dependent cell protrusion dynamics and traction force fluctuations in these conditions were plotted in Figure 1d. Here we used the grouped-clutch algorithm to significantly enhance the computational efficiency by grouping clutches together to have a smaller number of clutches to represent all clutches, which produced equivalent results but with much faster simulations (Figure S2). In Condition I, a typical simulated migrating cell with balanced motor and clutch number showed the dynamic protrusions with sequential phases of nucleation, elongation, retraction, and elimination, and fluctuating traction force to produce fast cell migration (Figure 1c,1d, Condition I). In Condition II with the higher motor number in the cells, the motor-clutch mechanism became “free-flowing” [21] with faster F-actin flow, lower traction force, shorter protrusion length, and hence slower cell migration (Figure 1c,1d, Condition II). In Condition III with the higher clutch number, the motor-clutch system became “stalled” [21] with near zero F-actin flow, near-maximal constant traction force, limited dynamic protrusions, and hence poor cell migration (Figure 1c,1d, Condition III). In Condition IV with the higher actin polymerization rate, the protrusion dynamics became more significant, with longer protrusion length and highly fluctuating traction forces, to produce faster cell migration (Figure 1c,1d, Condition IV). Overall these simulations indicate that the CMS fundamental parameters (*n*_m_, *n*_c_, *v*_poly_) can uniquely govern cell migration.

### Heterogeneity in migration phenotypes of glioblastoma patient cells

To understand the migration phenotypes of glioblastoma patient cells, we measured the cell migration of Mayo glioblastoma PDX lines of MES (MM, 3 lines), PN (MP, 3 lines), CL (MC, 5 lines) subtypes with adherent culture [34] and UCSD glioblastoma PD lines of MES (UM, 1 line), PN (UP, 1 line) subtypes cultured as neurospheres [35], for a total of N=13 and lack of correlation between motility, traction strain energy, and F-actin flow rate patients (Figure 2c). We measured the migration of glioblastoma cells on PA gels with different stiffnesses, coated with laminin for the Mayo cells and Matrigel for the UCSD cells, to reach adequate cell adhesion (Figure 2a,2b).

**Figure 2.**
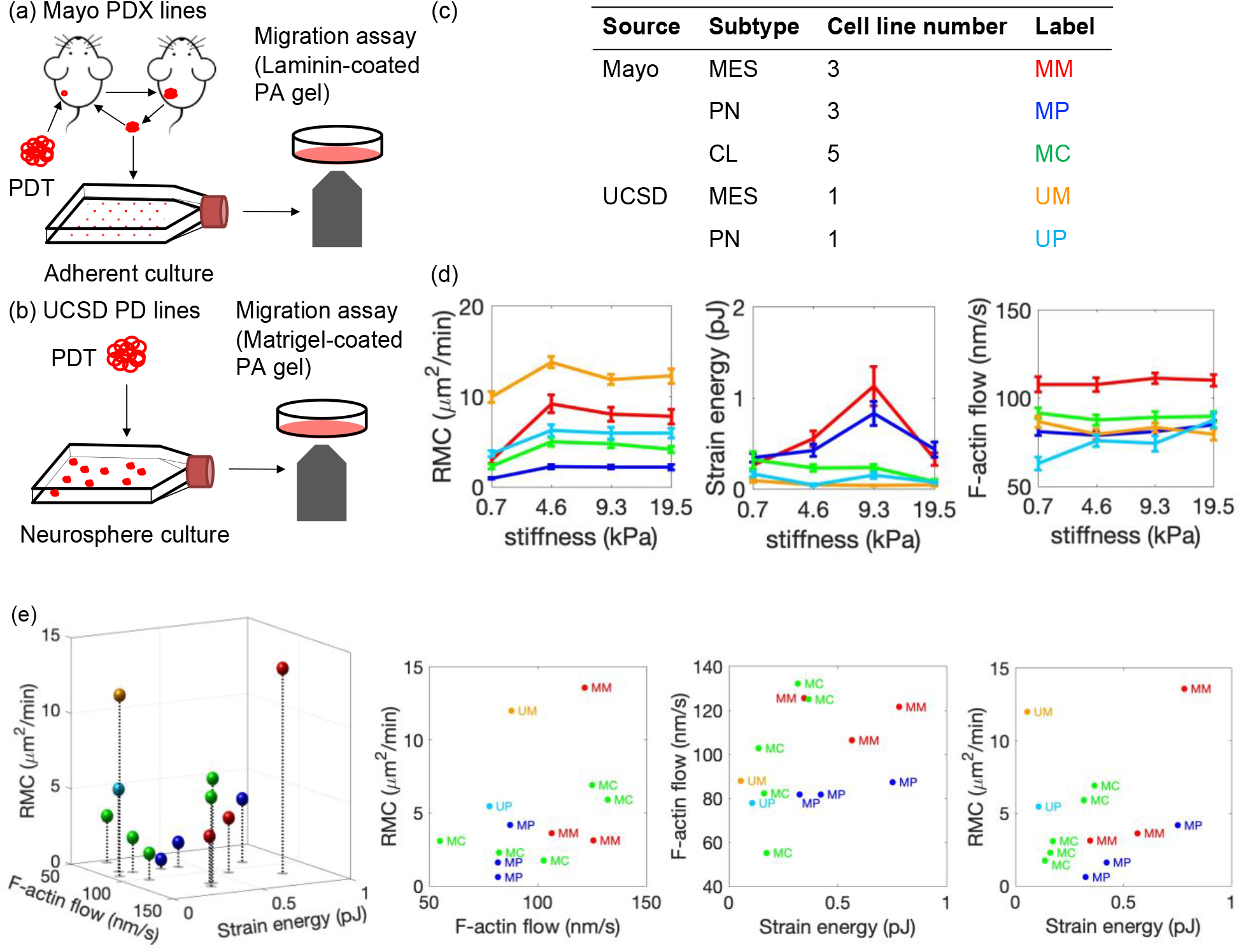
Heterogeneity in migration phenotypes of glioblastoma patient cells. (a) Mayo PDX cell lines were established as previously described by implanting patient tumors into mouse flanks [34], and then cultured in adherent conditions with FBS media. (b) UCSD PD cell lines were established as described previously [35], and cultured in neurosphere conditions with NSC Media. Mayo and UCSD cells were plated on polyacrylamide gels with Young’s modulus of 0.7, 4.6, 9.3, 19.5 kPa, coated with laminin and Matrigel, respectively. Cell motility (random motility coefficient, RMC), traction strain energy, and F-actin retrograde flow were measured using established protocols [26]. (c) There were 3 Mayo MES lines (MM), 3 Mayo PN lines (MP), 5 Mayo CL lines (MC), 1 UCSD MES line (UM), and 1 UCSD PN line (UP). (d) The mean ± SEM values of RMC, strain energy, F-actin flow of Mayo MES, PN, CL cells and UCSD MES, PN cells on PA gels with different stiffnesses were highly variable across cell lines. (e) The mean values of RMC, strain energy, F-actin flow combining all substrate stiffnesses for each cell line were plotted in the 3D space, along with their 2D projections. RMC, tractions strain energy, and F-actin flow were all highly variable with no obvious correlations with each other.

We found heterogeneity in cell migration of glioblastoma cells with different subtypes and sources, and their mean ± SEM of cell motility (RMC), traction strain energy, and F-actin retrograde flow rate of glioblastoma cells on PA gels with different stiffnesses were all highly variable (Figure 2d). Despite this heterogeneity, we found that glioblastoma cells tended to have maximal motility on stiffnesses ranging from 4.6 kPa to 19.5 kPa (Figure 2d), which is closer to brain tissue stiffnesses (1 – 6 kPa [36,37]). Mayo MES and PN cells exhibited optimal traction strain energy with stiffness of 9.3 kPa, while the other cell lines had low cell traction strain energy with no evidence of optimality (Figure 2d). All cell lines had no clear optimal F-actin retrograde flow (Figure 2d), cell area, and aspect ratio (Figure S3b) as a function of substrate stiffness. The differences in cell migration between subtypes and sources were similar at all substrate stiffnesses (i.e., Mayo MES had higher motility than Mayo PN cells at all stiffnesses) (Figure 2d), and therefore in subsequent analysis we combined the cell migration data for a given PD(X) line across all substrate stiffnesses.

The mean cell motility (RMC), traction strain energy, and F-actin flow rate for each PD(X) line was plotted in the 3D experimental measurement space with 2D projections defined by these three measurables (Figure 2e). The cell migration data of all glioblastoma cells across all subtypes and sources was plotted in Figure S3c with statistical analysis. Mayo MES cells had significantly higher motility, F-actin flow, cell area and aspect ratio compared to Mayo PN cells (Figure 2e, S3c). Mayo CL cells had intermediate values in motility, F-actin flow, and morphology, except for the lower traction strain energy, compared to Mayo MES, PN cells (Figure 2e, S3c). UCSD MES cells had higher motility and cell area compared to UCSD PN cells (Figure 2e, S3c). All Mayo cells had higher traction strain energy, F-actin flow, and cell area, with lower motility and aspect ratio compared to all UCSD cells (Figure 2e, S3c). Overall, these results show significant heterogeneity in cell migration mechanics of glioblastoma cells across different subtypes and sources, and general lack of correlation between motility, traction force, and F-actin flow rate.

### CMS-derived physical parameters of glioblastoma patient cell migration

To transform these empirical measurements of glioblastoma patient cell migration mechanics into fundamental mechanistic interpretation, we used the CMS to parameterize the cell migration of PD(X) lines by fitting their motility, traction force, and F-actin retrograde flow experimental data to simulations with adjusted physical parameters (*n*_m_, *n*_c_, *v*_poly_). For each PD(X) line this effectively mapped the 3D empirical observation space into a 3D theoretical physical space. These physical parameter values were then plotted in the 3D CMS parameter space of (*n*_m_, *n*_c_, *v*_poly_) for each patient with 2D projections in Figure 3a, and also plotted as bar graphs in Figure S4. We found that despite the apparent heterogeneity of physical parameters, glioblastoma PD(X) lines consistently showed approximately balanced motor number and clutch number (*n*_c_/*n*_m_~0.75) (Figure 3a), which enabled robust cell migration and traction forces that increased with the motor-clutch level (Figure 1c). Therefore, we can also plot the CMS parameter values on the heat map of various values for (*n*_m_, *v*_poly_) with a constant ratio *n*_c_/*n*_m_ = 0.75 in Figure 3b. We found that Mayo MES cells had higher motor (*n*_m_) and clutch (*n*_c_) number, and higher F-actin polymerization rate (*v*_poly_) compared to PN cells (Figure 3a,3b,S4), resulting in higher motility and F-actin flow. UCSD MES cells had higher F-actin polymerization rate (*v*_poly_) compared to UCSD PN cells, resulting in higher motility. Mayo cells had higher motor and clutch number and lower F-actin polymerization to produce higher traction force and lower motility compared to UCSD cells (Figure 3a,3b). These results suggested that myosin motors and adhesion clutches were well balanced in all glioblastoma cells, that fast-moving cells had higher F-actin polymerization with either high or low cell traction, and adherently cultured cells had higher myosin motors and adhesion clutches and lower F-actin polymerization compared to neurosphere cultured cells. Overall, we find that the 3D CMS parameter space (*n*_m_, *n*_c_, *v*_poly_) provides a more fundamental and revealing framework for describing glioblastoma PD(X) migration than does the empirical 3D space defined by the measured experimental quantities (RMC, traction strain energy, F-actin retrograde flow).

**Figure 3.**
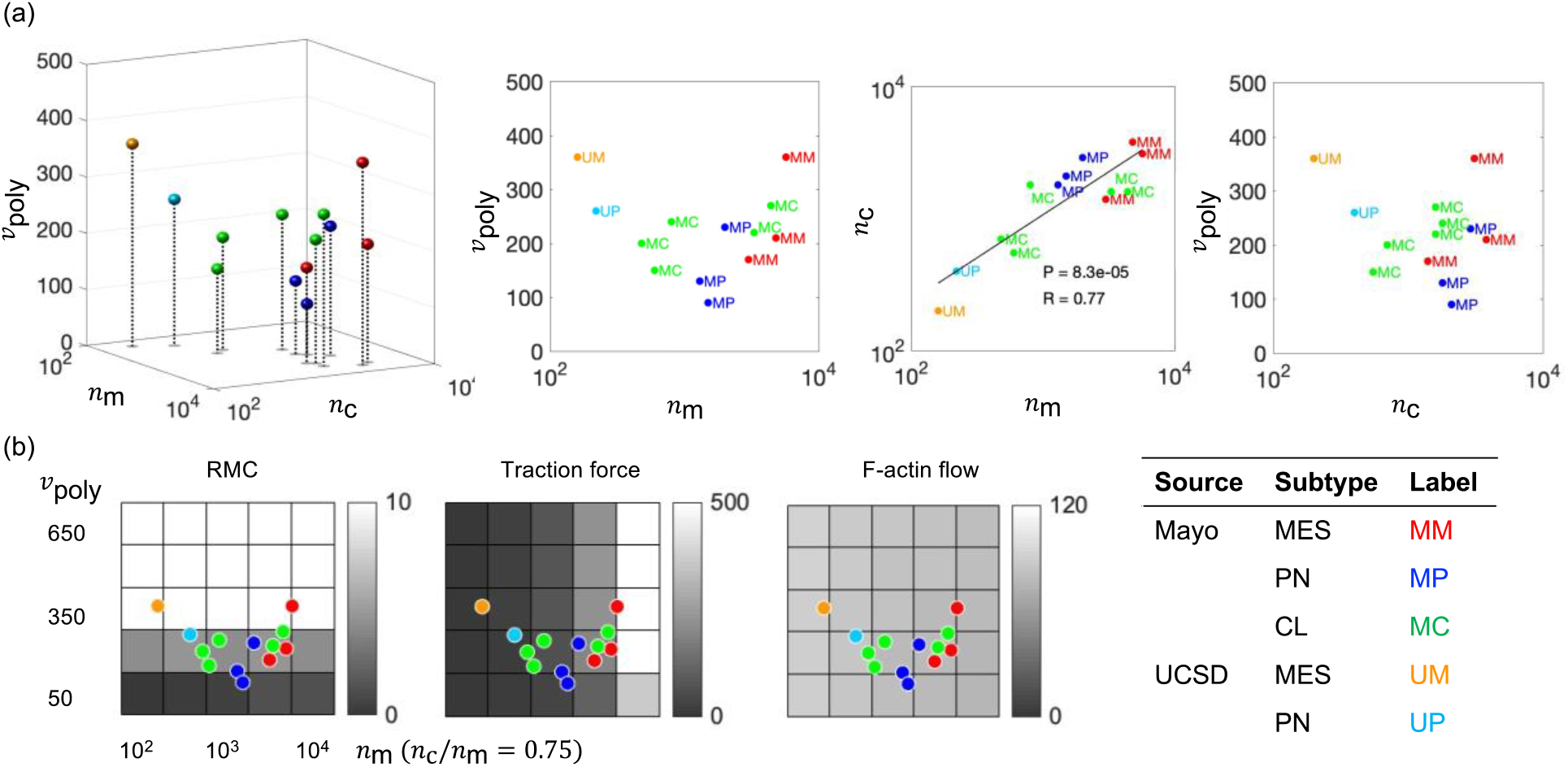
CMS-derived physical parameters of glioblastoma patient cell migration. (a) Physical parameter values of glioblastoma cells derived by fitting the cell migration with the CMS predictions and plotted in the 3D CMS parameter space of (*n*_m_, *n*_c_, *v*_poly_) with 2D projections. (b) Physical parameter values of glioblastoma cells were plotted in the heat map of the CMS prediction with various values for (*n*_m_, *v*_poly_) with a constant ratio between *n*_m_ and *n*_c_ (*n*_c_/*n*_m_ = 0.75). Given the balanced motor-clutch ratio, cell speed is largely determined by *v*_poly_.

### The CMS predicts PD(X) glioblastoma cell migration upon drug perturbations

Knowing the CMS 3D parameter set for each PD(X) line not only allows us to predict the cell migration using the CMS, but also to predict different migration behaviors with the change of parameter values due to drug treatments. The CMS predictions with various values of (*n*_m_, *v*_poly_) and a constant clutch number (*n*_c_ = 225) were plotted as the heat map in Figure 4a, along with the UCSD MES cells, which had higher F-actin polymerization rate (*v*_poly_) and lower motor number (*n*_m_) (Figure 4a orange) but higher motor/clutch ratio (Figure 3a), resulting in higher cell motility and higher F-actin flow (Figure 4a,4b orange), compared to UCSD PN cells (Figure 4a,4b blue).

**Figure 4.**
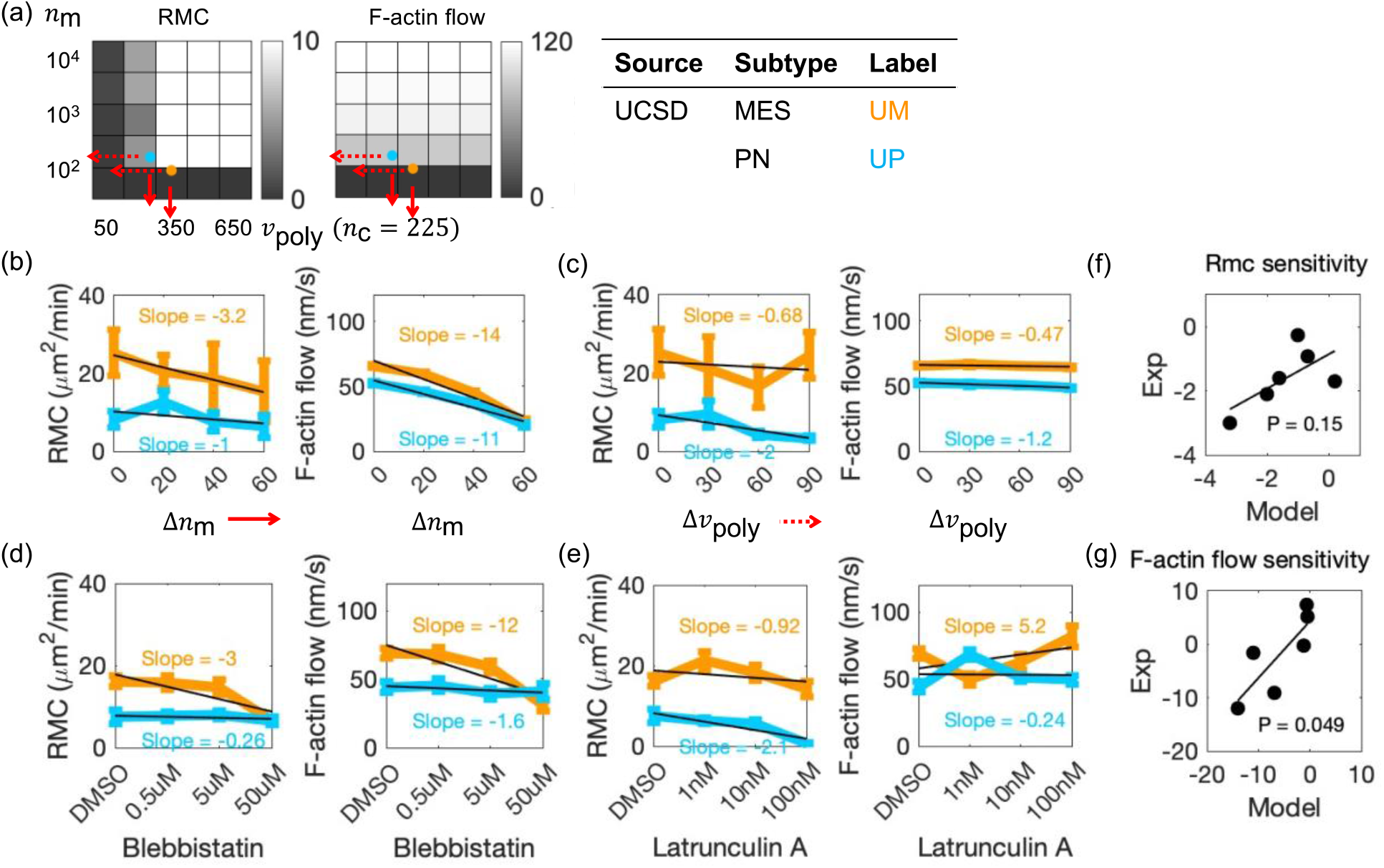
The CMS predicts differential cell migration and F-actin flow sensitivity upon actomyosin drug perturbation. (a) The heat map of the CMS prediction, including motility (RMC) and F-actin retrograde flow, with various values of (*n*_m_, *v*_poly_) with a constant value of *n*_c_ = 225. UCSD MES cells (orange) had a higher F-actin polymerization rate and lower motor number compared to UCSD PN cells (light blue), which produced higher motility and higher sensitivity of F-actin flow with reducing the motor number. (b,c) The mean ± SEM values of RMC and F-actin flow of UCSD MES and PN cells with the decreasing motor number (b) (solid-red arrow) and decreasing F-actin polymerization rate (c) (dotted-red arrow) predicted by the CMS were plotted. (d,e) The mean ± SEM values of RMC and F-actin flow of UCSD MES and PN cells with different concentrations of myosin II inhibitor, blebbistatin (d), and F-actin polymerization inhibitor, Latrunculin A (e), were plotted. The RMC (f) and F-actin flow (g) sensitivities to decreasing parameter values predicted by the CMS (x-axis in (f,g), also slopes in (b,c) and Figure S5 (b,c)), were correlated with the sensitivities to cytoskeletal drugs (y-axis in (f,g), also slopes in (d,e) and Figure S5 (d,e)), with p = 0.15 and p = 0.05, respectively.

When reducing the motor number, we estimated that the motility and F-actin flow of UCSD MES cells would be reduced to a greater extent than UCSD PN cells as indicated by the solid red arrows in Figure 4a, which was confirmed by the CMS predictions with the reducing motor number (Δ*n*_m_) in Figure 4b. Consistent with the CMS predictions, when UCSD cells were treated with blebbistatin to inhibit their myosin II contractility, the cell motility and F-actin flow of UCSD MES cells on PA gels decreased more significantly compared to UCSD PN cells (Figure 4d). This test of the model was not only consistent with the CMS predictions (Figure 4b), but also confirmed the motor number difference between UCSD MES and PN cells (Figure 3a,4a).

When reducing the F-actin polymerization rate, we predicted that the UCSD PN cells would reduce their motility to a greater extent than UCSD MES cells, with the insensitivity of Factin flow, as indicated by the dotted arrow in Figure 4a, also confirmed by the CMS predictions with the reducing F-actin polymerization rate, Δ*v*_poly_, in Figure 4c. We then treated the UCSD cells with Latrunculin A to inhibit their actin polymerization by binding G-actin monomers, and found the motility of UCSD PN cells on PA gels indeed decreased more significantly compared to UCSD MES cells (Figure 4e), which again was consistent with the CMS predictions (Figure 4c).

Adherent Mayo cells had higher motor and clutch number and lower F-actin polymerization rate compared to neurosphere UCSD cells in Figure 3. To test whether the culture conditions affect the migration phenotype, we cultured the neurosphere UCSD MES cells adherently for one week to create UCSD MES-AD cells before conducting migration assays, and we found the migration phenotypes of UCSD MES-AD cells became closer to the phenotypes of UCSD PN cells, with higher motor number and lower F-actin polymerization rate resulting in lower motility and higher F-actin flow compared to UCSD MES cells (Figure S5a, S5b). We also treated the UCSD MES-AD cells with cytoskeleton drugs, and found MES-AD cells had lower sensitivity to blebbistatin and higher sensitivity to Latrunculin A in motility and F-actin flow compared to UCSD MES cells (Figure S5d, S5e), which again was consistent with the CMS predictions (Figure S5b, S5c), and confirmed that adherent culture can increase the functional motor number and decrease the F-actin polymerization rate of neurosphere cultured cells (Figure S5a)

Overall, we found that CMS-predicted sensitivities of cell motility (Figure 4f) and F-actin flow (Figure 4g) to decreasing parameter values were highly correlated with the measured sensitivities to cytoskeletal drugs. Not only can we use the CMS physical parameter values of glioblastoma PD(X) lines to not only describe migration phenotypes (Figure 3), but also to predict the differential migration changes between patients due to the changes in the parameter values either by cytoskeletal drugs or by the culture conditions (Figure 4, S5).

### CMS-based transcriptomic biomarkers for glioblastoma cell migration

In principle, the CMS parameters should depend on transcriptional levels of key genes controlling motor, clutch, and F-actin polymerization activities. Thus, we sought to identify correlates of the CMS parameters in previously collected transcription level data. In Figure 3 and S4, there are consistent differential estimates of the CMS parameters between MES and PN for both Mayo and UCSD patient cells. Moreover, PN and MES subtypes became more frequent in patients with tumor recurrences [15]. Therefore, we decided to analyze the differential genes between MES and PN and correlated them with the CMS parameters. We first derived RNAseq RPKM expression of Mayo PDX cells from Vaubel et al., 2020 [34] (19552 genes in 20 MES patients, 16 PN patients). We filtered out genes with small mean values and low patient portions to achieve a normal gene expression distribution (Figure S6a, S6b, 11,752 genes left). We derived 1,177 differential genes between Mayo MES and PN cells using the two-sample t-test (p<0.05, FC>2, Figure S6c). We applied enrichment analysis to these differential genes in Gene Ontology [38] and derived 216 migration genes in the “cell adhesion” and “cell motility” biological processes. We further applied the enrichment analysis to these migration genes and derived 23 actin genes, 9 motor genes, and 47 clutch genes in the “actin cytoskeleton,” “actomyosin” coupled with “myosin II complex,” and “focal adhesion” cellular components, respectively. This resulted in a final reduced list of actin, motor, and clutch genes that could potentially be correlated with the CMS physical parameters.

We applied correlation analysis between the mRNA expression of the actin, motor, clutch genes in the 10 Mayo PDX lines used in the present study for which mRNA expression level was available, and their CMS parameter values (*v*_poly_, *n*_m_, *n*_c_), respectively. In the actin genes, VCL had the highest positive correlation coefficient (R) and ADD2 had the lowest R, with the actin polymerization rate (Figure 5). However, none of the correlations were statistically significant at the p=0.05 level. Among the motor genes, FBLIM1, ACTN1, PALLD, and MYL12A had statistically significant and positive R (p<0.05), while PRKCZ had the lowest negative R with the motor number, but it was not statistically significant at the p=0.05 level (Figure 5). In the clutch genes, FBLIM1, VCL, CSRP1, CD44, ITGB1, MSN, and CD99 had positive R that was statistically significant (p<0.05), and ACTN2 had the lowest R with the clutch number but it was not statistically significant at the p=0.05 level (Figure 5). Thus, there is a subset of genes expressed that were highly correlated with the motor and clutch CMS parameter values, and these were all in the direction expected, i.e., positive correlations imply that higher gene expression levels correlate with higher functional number of motors and clutches. Using the 95% confidence interval we would expect that only 5% of the 79 total genes, or ~4 genes, would have statistical significance by chance alone, and their slopes would be a random mixture of positive and negative correlations. Instead, we observed that 11 genes were significantly correlated (p<0.001 for a Poisson distribution with mean=4), and all were positively correlated (1/2^11^=0.0005). All of the 11 genes were associated with either motor or cutch values, while none were associated with F-actin polymerization activities. We also applied Cox regression analysis between the mRNA expression ratios of the 79 migration genes and Mayo patient overall survival (OS), and we found 3 genes (ALCAM, SDC4, ITGB5) had significant and positive hazard ratios of the OS (Figure S7), which was consistent with the significance by chance (5% of 79 genes is ~ 4 genes). Overall, our CMS parameters define mechanical biomarkers that can potentially be converted into transcriptomic biomarkers that can be used to predict the cell migration of glioblastoma patient cells.

**Figure 5.**
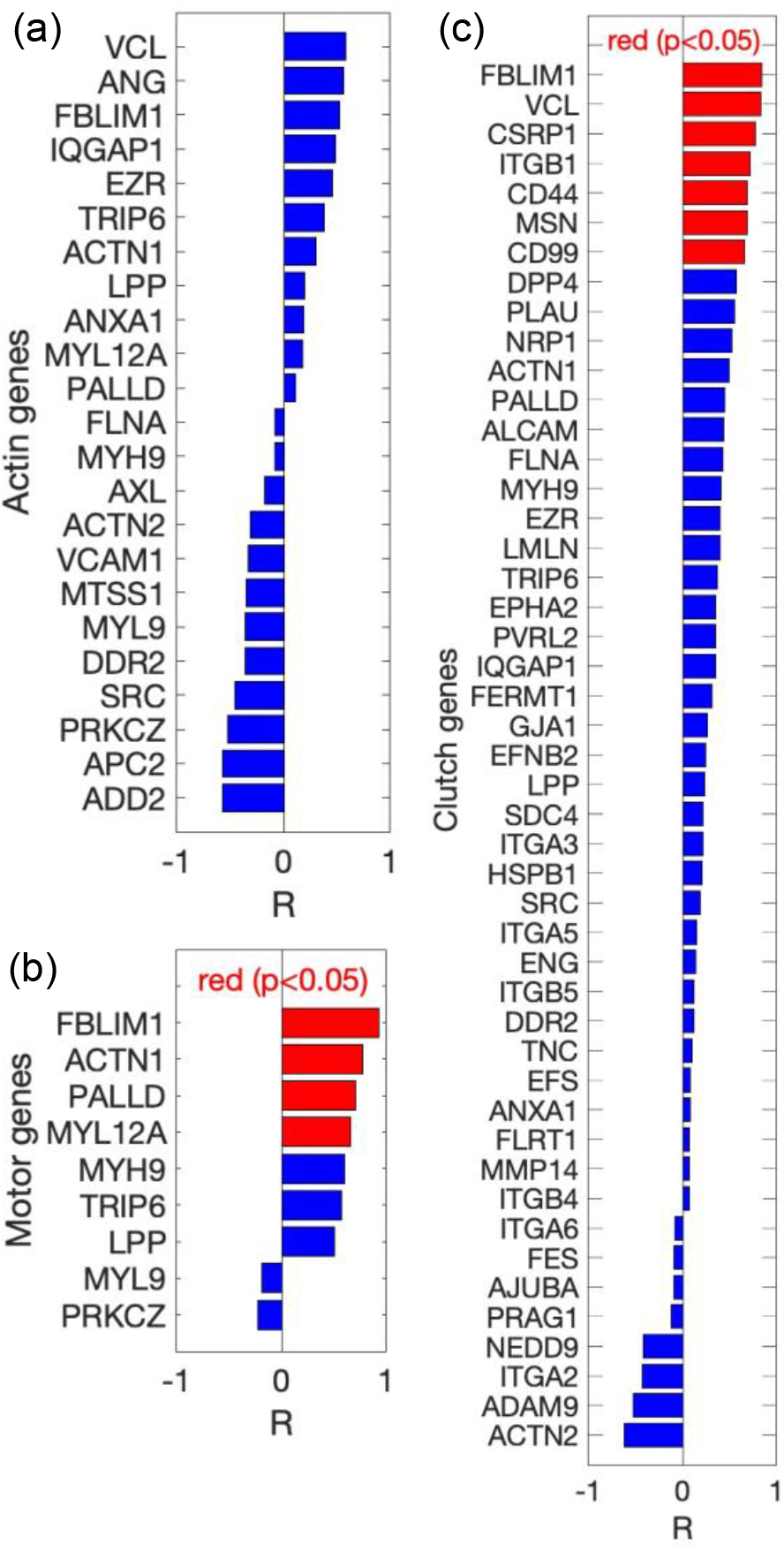
Identification of CMS-based transcriptomic biomarkers for glioblastoma cell migration. We derived 23 actin genes (a), 9 motor genes (b), and 47 clutch genes (c) based on Gene Ontology cellular components from 1,177 differential genes between Mayo MES and PN cells (Method: mRNA expression analysis). We applied the correlation analysis between the mRNA expression ratios of the actin (a), motor (b), clutch (c) genes in the 10 Mayo lines used in the present study (no mRNA data was available for the Mayo 16 line) and their CMS parameter values, *v*_poly_ (a), *n*_m_ (b), *n*_c_ (c), respectively, along with their correlation coefficients (R) were sorted and plotted in (a-c) (significant correlation is in red).

## Discussion

In this study, we used the physics-based motor-clutch model, termed here the CMS [26] (Figure 1a), to mechanistically parameterize and predict glioblastoma cell migration mechanics and speed. We reduced the 11-dimensional parameter space of the CMS into 3 dimensions based on their parameter sensitives (Figure 1b), and identified three fundamental physical parameters: motor number, clutch number, and actin polymerization rate that can uniquely govern cell migration (Figure 1c,1d). We found significant heterogeneity in glioblastoma patient cell migration across subtypes and sources (Figure 2) and derived the physical parameter values for each cell line by fitting their cell migration with the CMS predictions (Figure 3). Despite their heterogeneity, glioblastoma cells had balanced motor/clutch ratios (*n*_c_/*n*_m_ ~0.75) to produce robust cell migration and traction force (Figure 1c,3a). In addition, we found consistent trends by molecular subtype, with Mayo MES cells having higher motor-clutch number and F-actin polymerization rate relative to Mayo PN cells (Figure 3), resulting in higher motility and F-actin flow (Figure 2e). Similarly, UCSD MES cells had higher F-actin polymerization rate relative to the UCSD PN cells (Figure 3) resulting in faster migration (Figure 2e). Moreover, the CMS accurately predicted the differential sensitivities between MES and PN cells to cytoskeletal drug perturbations in cell motility and Factin flow according to their differential physical parameters. Finally, we derived a list of motor-clutch-associated genes in the Mayo cells having mRNA expression correlating with the physical parameters of Mayo cells, which can potentially be used to predict CMS cell migration parameters and speeds. Overall, we describe a simplified 3D physics-based framework for mechanically parameterizing individual glioblastoma patients and connecting biomechanics to clinical transcriptomic data, which can potentially be used to predict the cell migration and drug responses in glioblastoma cells.

We found that glioblastoma MES cells have faster migration than PN and CL cells on compliant PA gels coated with different extracellular matrices (ECMs), which is consistent with Munthe et al., 2016 [12] suggesting that glioblastoma MES cells had higher motility compared to PN and CL cells based on the collective cell migration of free-floating spheroids. Piao et al., 2013 [39] similarly found the glioblastoma cell lines resistant to bevacizumab were similar to the MES subtype and had higher invasive capacity and motility compared to other subtypes. To provide the biophysical mechanisms of the differential cell motility, we parameterized the cell migration of glioblastoma cells with the CMS and found the higher F-actin polymerization rate best explained the faster migration of the MES cells relative to the PN cells (Figure 3, S4). In the CMS simulation, higher F-actin polymerization rate promotes cell protrusion dynamics, with longer protrusion length and highly fluctuating traction force, causing highly unbalanced protrusion forces, highly polarized cell morphology, and hence faster migration (Figure 1c, 1d). The CMS also predicted the higher sensitivities of UCSD PN cells to the actin-inhibiting drug, Latrunculin A, due to the lower F-actin polymerization rate relative to MES cells (Figure 4a, 4c, 4e). Adebowale et al., 2021 [30] similarly confirmed the significant filopodia dynamics with longer filopodia length and lifetime resulted in faster cell migration of fibrosarcoma cells on fast-relaxing viscoelastic gels, demonstrating the protrusion dynamics promoting cell motility *in vitro*, which was also supported by the CMS simulation coupled with viscoelastic substrate. Therefore, we find that F-actin polymerization becomes a potential target to alter glioblastoma cell motility, and the CMS can potentially predict the patient-specific cell motility and treatment responses.

It is interesting that the CMS parameterized the glioblastoma patient cell migration to derive balanced motor and clutch number (*n*_c_/*n*_m_ ~0.75) (Figure 3a), which produces robust cell migration and traction forces (Figure 1c, 1d) in the CMS simulation. The balanced motor and clutch number can be explained by protein interactions between myosin II and focal adhesions. In cell migration, myosin II mediates focal adhesion formation and maturation by bundling actin filaments, clustering adhesion proteins, and inducing conformational changes to promote adhesion protein binding [27]. Focal adhesions also activate the RHOA and ROCK signaling and further activate myosin II and actomyosin contraction [27]. Therefore the positive association between motor and clutch also manifests itself in different cell types, such as the large focal adhesions and high myosin II activation resulting in higher traction forces in fibroblasts, compared to the high motility and low traction force with smaller adhesion size in leukocytes [27,28]. By inhibiting myosin II with blebbistatin and inhibiting integrin-mediated adhesion with cyclo(RGCfV) for U251 cells, we can maintain the cell motility but significantly reduce the traction force [26]. In this study, we also show that adherently cultured Mayo cells have much higher motor-clutch number resulting in higher traction force than neurosphere-cultured UCSD cells (Figure 2,3), showing that culture condition can significantly impact patient cell migration. By culturing UCSD MES cells adherently for a week to become MES-AD cells, we further increased the motor-clutch number and decreased the F-actin polymerization rate, resulting in higher F-actin flow and lower motility with lower sensitivity to myosin II inhibitor, blebbistatin, consistent with the CMS simulation (Figure S5). We also note that the optimality of cell motility and traction force as a function of stiffness, reported previously for U251 cells in vitro (Bangasser et al., 2017), is also evident in many of the PD(X) cell lines. Overall, we find that the interactions between myosin II and focal adhesions support a balanced motor-clutch number in glioblastoma cells, and the CMS can potentially predict the cell migration and treatment responses based on patient-specific motor-clutch ratio.

To find the correlated genes with the CMS parameters, we analyzed the differential genes between Mayo MES and PN cells (20 MES patients, 16 PN patients). We identified 23 actin genes, 9 motor genes, and 47 clutch genes based on the cellular components in Gene Ontology, and we found 11 genes were significantly correlated with the CMS parameters (Figure 5), all of which were positive correlations (11 correlated genes are substantially higher than the ~4 expected for randomly correlated genes in 79 total genes with p=0.05), showing that the CMS parameters closely and positively correlated with genes in myosin II and focal adhesion cellular components (Gene Ontology). Migfilin (FBLIM1) is closely associated with myosin II activation [40], α-actinin (ACTN1) cross-links F-actin filament and collaborates with actomyosin system [41], palladin (PALLD) binds F-actin and α-actinin to modulate actomyosin network [42], myosin light chain (MYL12A) colocalizes with actin stress fibers and is the key component in myosin II [43], and suggesting possible mechanisms for these genes to correlate with motor number. Migfilin (FBLIM1) regulates integrin activation [40], vinculin (VCL) is a focal adhesion protein that binds F-actin and talin [44], cysteine-rich protein (CSRP1) interacts with adhesion proteins zyxin and α-actinin [45], integrin-β1 (ITGB1) is a fundamental focal adhesion receptor that spans the plasma membrane and links the ECM to F-actin system [46], CD44 is a plasma membrane-spanning glycoprotein that binds hyaluronan in the ECM [47], moesin (MSN) binds F-actin and CD44 to regulate focal adhesions [48], CD99 is a transmembrane protein that activates cell adhesion protein ICAM1 and FAK to regulate focal adhesions [49]. Overall, these genes are closely associated with the CMS parameters, and are potential markers and targets to predict and interrupt the glioblastoma cell migration based on patient-specific transcriptomic information.

Our study has some limitations and points to future work. While we would expect that the faster migration speed in MES cells may contribute to the lower survival found in MES patients vs. CL or PN [14,15], we suspect that other confounding variables need to be included in the further analysis, such as age [3] and immune response [14,15] in order to better predict patient survival. While we had hoped to identify genes that associate with F-actin polymerization rate, which contributes to the faster migration in MES cells, we failed to find any statistically significant correlations. It is likely that we need a larger cohorts of patient cells (only 3 MES, 3PN, and 5 CL Mayo PDX lines in this study) with greater sequencing depth in actin, motor, and clutch genes. In order to be of clinical utility, future work will need to prospectively test transcriptomic predictions in terms of cell migration and drug sensitivities, rather than the retrospective relationships identified in the present study. Even so, overall the results provide a proof-of-concept that CMS-based mechanical biomarkers can be used to describe cell migration dynamics, predict differential drug sensitivities, and identify correlations with mechanistically relevant mRNA transcript levels.

## Materials and Methods

### Cell Migration Simulator (CMS)

The detailed governing equations and algorithms of the Cell Migration Simulator (CMS) was described in Bangasser et al., 2017 [26]. Briefly, the CMS comprises multiple protrusions or modules which were nucleated randomly based on the rate *k*_mod_ which is a function of the maximum rate 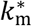, G-actin (*A*_G_) or free actin monomers, and the total actin amount (*A*_T_) described in Equation 1.

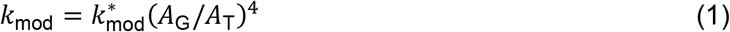

Protrusions were elongated based on the polymerization rate *v*_poly_ as a function of the maximum rate 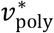, *A*_G_, and *A*_T_ written in Equation 2.

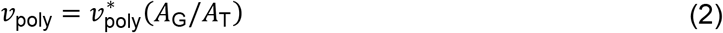

Protrusions were capped randomly at the rate *k*_cap_ eliminating further polymerization and removed if the protrusion length is shorter than the minimum length *l*_min_.

The cell position was determined by the force balance between protrusion forces *F_j_* for *n*_mod_ modules and the cell body force *F*_cell_ based on Equation 3.

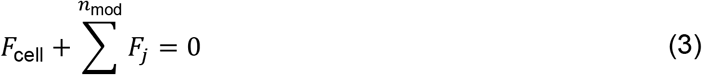

*F*_cell_ and *F_j_* were determined based on the motor-clutch model [20]. *F*_cell_ and *F_j_* are summation of individual clutch force *F*_c, *i*_, which is a function of the clutch spring constant κ_c_ and clutch displacements *dx*_c_, in Equation 4.

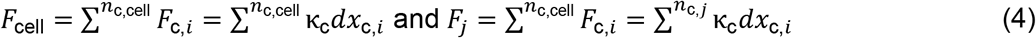

Clutches bound and unbound to F-actin based on the clutch binding rate *k*_on_ and unbinding rate *k*_off_ as a function of the minimum unbinding rate 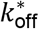, motor force *F*_m_ and *F_j_*, and *F*_m_ equals to the single motor force 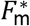 multiplying the motor number *n*_m_ in Equation 5.

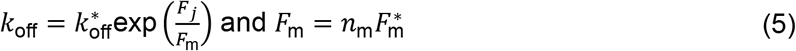

Bound clutches were displaced at the rate *v*_actin_ as a function of the maximum rate 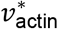, *n*_m_, *F*_m_, and *F_j_* in Equation 6.

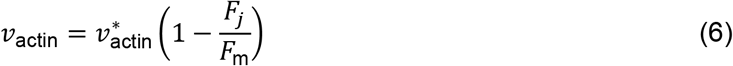

Monte Carlo simulations were conducted using a direct Gillespie Stochastic Simulation Algorithm, the event was executed based on accumulated event rates, including *k*_on_, *k*_off_, *k*_mod_, and *k*_cap_, and the next time step *t*_step_ was determined based on the total event rates ∑*k_i_* and a random number *RN* in Equation 7.

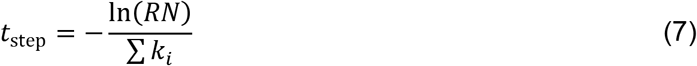

The C++ version of the CMS [29] was used to conduct the simulations in the Mesabi computer cluster at the Minnesota Supercomputing Institute (MSI).

### Grouped clutch

In the CMS simulations, we found that computational time increased proportionally with the clutch number, since the larger clutch number caused the smaller time step and hence longer computational time based on the direct Gillespie Stochastic Simulation Algorithm (SSA) [26]. In order to enhance the computational efficiency, we grouped multiple clutches into one clutch with the summed clutch stiffness in the CMS, by assuming that multiple clutches had similar reactions during adhesion forming (Figure S2). In the CMS governing equations, the module force and the cell body force are the summation of each clutch deformation multiplying the clutch spring constant, as in Equation 4. With *g* clutches were grouped, the force equations were rewritten as in Equation 8.

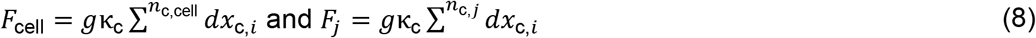

We found the CMS with grouped clutches had equivalent predictions compared to the CMS with individual clutches, showing that with enough clutch number, larger clutch number can be described by smaller clutch number, and each clutch representing a group of clutches contributed a similar mechanical force to the system, compared to the force sum of the clutch group, and hence the system remained equivalent (Figure S2). Therefore, with prescribed large clutch number, such as 10000, the system can be simulated as with 750 clutches, which dramatically reduced computational time. In this study, we used the CMS with grouped clutches to conduct simulations, which reduced the computation time more than 10-fold.

### Parameter sensitivity and clustering

The cell migration predictions, including the maximum RMC, the maximum traction force, and the minimum F-actin flow over different stiffnesses were generated by the CMS with the changes of the base parameter values (Table 1), plotted in Figure S1. The linear regression between the CMS migration predictions Y and the logarithm of parameter ratios were plotted in Figure S1. Parameter sensitivity values were determined by the slope of the linear regression normalized by the base prediction values (Y_0_) (slope (linear regression) / Y_0_). We applied the agglomerative hierarchical clustering to the CMS parameter sensitivities using the linkage function in MATLAB with average method to identify the main clusters for the CMS parameter sensitivities (Figure 1b).

### Glioblastoma patient cell lines and cell culture

Mayo PDX cell lines were developed and maintained by the Sarkaria lab at Mayo Clinic (Rochester, MN) (ref.). Cell lines were established by implanting patient tumors into mouse flanks, and cells were derived in short-term explant cultures with serum-containing medium [34,50]. We used MES (Mayo 16, 46, 59), PN (Mayo 64, 80, 85), CL (Mayo 6, 38, 76, 91, 195) cells. Cells were shipped in DMEM + 10% serum and grown adherently until confluent, and then frozen in 10% DMSO 90% FBS solution. Cells prepared for experiments were thawed into a flask coated with 10% GF-ReducedMatrigel (Corning 354263) in Neural Stem Cell (NSC) Media (DMEM/F12 (Gibco 11320033) + 1X B-27 Supplement (Gibco 12587010) + 1X Pen/Strep (Corning 45000-650) + 1ng/mL EGF/FGF (Peprotech AF10015 / Peprotech AF10018B) (added every 2-3 days)). Cells were allowed to recover for several days prior to imaging.

UCSD PD lines were developed and maintained by Clark Chen Lab at UMN (formerly of UCSD; ref.). Cell lines were derived and established from MES and PN glioblastoma patients, and cultured as neurospheres [35,51]. UCSD cells prepared for experiments were propagated in ultra-low adhesion flasks (Corning 3814) with NSC Media and were allowed to recover thawing for several days prior to imaging. For adherent culture conditions, UCSD cells were grown on a GF-Reduced Matrigel coated T-flask in NSC Medium for a minimum of 1 week prior to imaging.

### Preparation of ECM-coated polyacrylamide gels

Polyacrylamide (PA) gels with different stiffnesses (0.7kPa, 4.6kPa, 9.3kPa, 19.5kPa) were synthesized following the previous protocols [26]. In short, prepolymer mixtures of 40% acrylamide solution (Fisher BP1402), 2% bis-acrylamide solution (Fisher BP1404), 1 M HEPES (Sigma H6147) solution, and deionized water were prepared for the desired Young’s moduli, with 1% (v/v) 200 nm crimson fluorospheres (Invitrogen F8806) added for cell traction experiments. Mixtures were then polymerized by adding 0.6% (v/v) 1% ammonium persulfate (Bio-Rad 161-0700) solution and 0.4% (v/v) TEMED (Fisher BP150), followed by being quicky pipetted onto salinized glass culture dishes (MatTek P35G-0-20-C) and covered with a circular cover slip (Fisher 12-545-80) to produce ~100 μm thick gels. After polymerization, PA gels were activated with 0.5 mg ml^−1^ sulfo-SANPAH (Thermo 22589) and cultured with desired ECM molecules to conjugate these ECM molecules.

Mayo cells were plated on laminin with media containing 2% serum to promote adhesion. UCSD cells could not adhere to laminin, collagen, fibronectin, except for Matrigel with no serum.

### Live cell imaging

Time-lapse light microscopic images were taken using established protocols [26]. In short, glioblastoma cells were plated on PA gels overnight before imaging, Mayo cells were plated on laminin coated PA gels with media containing 2% serum to promote adhesion, and UCSD cells were plated on Matrigel coated PA gels without serum. Time-lapse phase-contrast images were taken for 10 h to track cell migration using a Nikon Eclipse TE200 microscope with a Plan Fluor 10 *×* /0.30NA objective. Time-lapse phase-contrast images were taken for 3 min to measure Factin flow using a Nikon Eclipse TE200 microscope with a Plan Fluor ELWD 40 × /0.60NA objective. Phase-contrast and epifluorescence images were taken before and after glioblastoma cells detached by treating with 0.05% trypsin to determine the cell traction using a Nikon Eclipse TE200 microscope with a Plan Fluor ELWD 40 × /0.60NA objective with a PhotoFluor II light source (89 North) and a mCherry/EGFP filter set.

Random motility coefficients (RMCs) of cell migration and cell area and aspect ratio were determined based on established protocols [26]. In short, cell position and region in 10-h timelapse images were tracked using a custom MATLAB code. The RMC of an individual cell was derived based on the mean squared displacement in the 2D diffusion equation described in Bangasser et al., 2017 [26]. The cell area and aspect ratio were calculated based on the recorded cell region. RMCs were measured for Mayo MES (Mayo 16(70), 46(90), 59(90)), PN (Mayo 64(100), 80(150), 85 (100)), CL(Mayo 6(50), 38(50), 76(50), 91 (50), 195 (30)) cells and UCSD MES (380), PN (180) cells on PA gels of 0.7, 4.6, 9.3, 19.5 kPa.

Actin retrograde flow of an individual cell was calculated based on established protocols [26]. Briefly, kymographs were made along the axis of moving protrusion features for 3-min time lapse images and actin flow was measured by analyzing the kymograph using a custom MATLAB code. Actin flow was measured for Mayo MES (Mayo 16(80), 46(50), 59(100)), PN (Mayo 64(130), 80(120), 85 (170)), CL(Mayo 6(80), 38(60), 76(60), 91(60), 195 (90)) cells and UCSD MES (250), PN (100) cells on PA gels of 0.7, 4.6, 9.3, 19.5 kPa.

Cell traction strain energy was determined using the established protocols based on the Fourier Transform Traction Cytometry [26]. Briefly, epifluorescence images before and after cells were detached were registered, aligned, and transformed using a custom MATLAB code. A gel displacement field was calculated by applying particle image velocimetry to the fluorospheres within the aligned epifluorescence images. The traction stress field was calculated using the Fourier Transform Traction Cytometry, and the total traction strain energy was calculated based on the dot products of the traction vectors and the displacement vectors. Strain energy was measured for Mayo MES (Mayo 16(40), 46(30), 59(40)), PN (Mayo 64(50), 80(30), 85 (50)), CL(Mayo 6(30), 38(30), 76(30), 91 (20), 195 (40)) cells and UCSD MES (100), PN (70) cells on PA gels of 0.7, 4.6, 9.3, 19.5 kPa.

### Parameterizing glioblastoma cell lines with (*n*_m_, *n*_c_, *v*_poly_) values

CMS predictions (*v*_F-actin_ (min), *F*_module_ (max), Rmc (max)) were linearly interpolated based on a three dimensional parameter space defined by *n*_m_, *n*_c_, and *v*_poly_. Cell traction force was estimated using the linear relation between module force and experimental strain energy (*F*_module_ = 175\**F*_strain_) based on the U251 maximum strain energy value and the assumption that U251 cells had 7500 clutches in the model [26]. The unique (*n*_m_, *n*_c_, *v*_poly_) values were found for each patient to limit the relative errors between CMS and experimental results to within 10% (For example, (*v*_F-actin_ (min, model) - *v*_F-actin_ (min, patient)) / *v*_F-actin_ (min, patient) < 10%).

### mRNA expression analysis

We derived RNAseq RPKM expressions of Mayo PDX cells from Vaubel et al., 2020 [34] (19,552 genes in 20 MES patients, 16 PN patients, and 30 CL patients, www.cbioportal.org). We filtered out genes with geometric means small than 1 and with counts in less than 80% of patients, and added a pseudo count of 0.1, to achieve a normal gene expression distribution compared to original distribution (Figure S6a,S6b, 11752 genes left). We applied the two-sample t-test to the RNAseq-derived mRNA expression levels of Mayo MES and PN lines, and derived 1,177 differential genes with p<0.05 and FC>2 in the volcano plot (Figure S6c). We applied enrichment analysis in Gene Ontology (http://geneontology.org) [38] and derived 216 migration genes in “cell adhesion” and “cell motility” biological processes. We further applied the enrichment analysis and derived 23 actin genes (Figure 5a), 9 motor genes (Figure 5b), and 47 clutch (Figure 5c) genes in “actin cytoskeleton,” “actomyosin” coupled with “myosin II complex,” and “focal adhesion” cellular components, respectively.

We applied the correlation analysis between the mRNA expression ratios of the actin (Figure 5a), motor (Figure 5b), clutch (Figure 5c) genes in the 10 Mayo lines used in the present study (no mRNA data available for Mayo 16 line) and their CMS parameter values (*v*_poly_, *n*_m_, *n*_c_), respectively. The correlation coefficients (R) of all the genes were sorted and plotted in Figure 5, with the significant correlation coefficients in red.

We applied Cox regression analysis between the mRNA expression ratios of the actin (Figure S7a), motor (Figure S7b), and clutch (Figure S7c) genes in a cohort of 66 Mayo patients and their overall survival, and their hazard ratios with 95% Confidence Interval were sorted and plotted in Figure S7, with the significant hazard ratios in red.

### Statistics

* Denotes p<.05, ** p<.01, *** p<.001 derived from the Kruskal-Wallis test with Dunn-Sidak *post hoc* analysis

## Acknowledgement

The study was supported by the National Institutes of Health via grants U54CA210190 (D.J.O.) and P01CA254849 (D.J.O.). The content is solely the responsibility of the authors and does not necessarily represent the official views of the National Institutes of Health.

## Code and data availability

All codes and data will be made available on reasonable request from the corresponding author.

## Supporting Information

**Figure S1.**
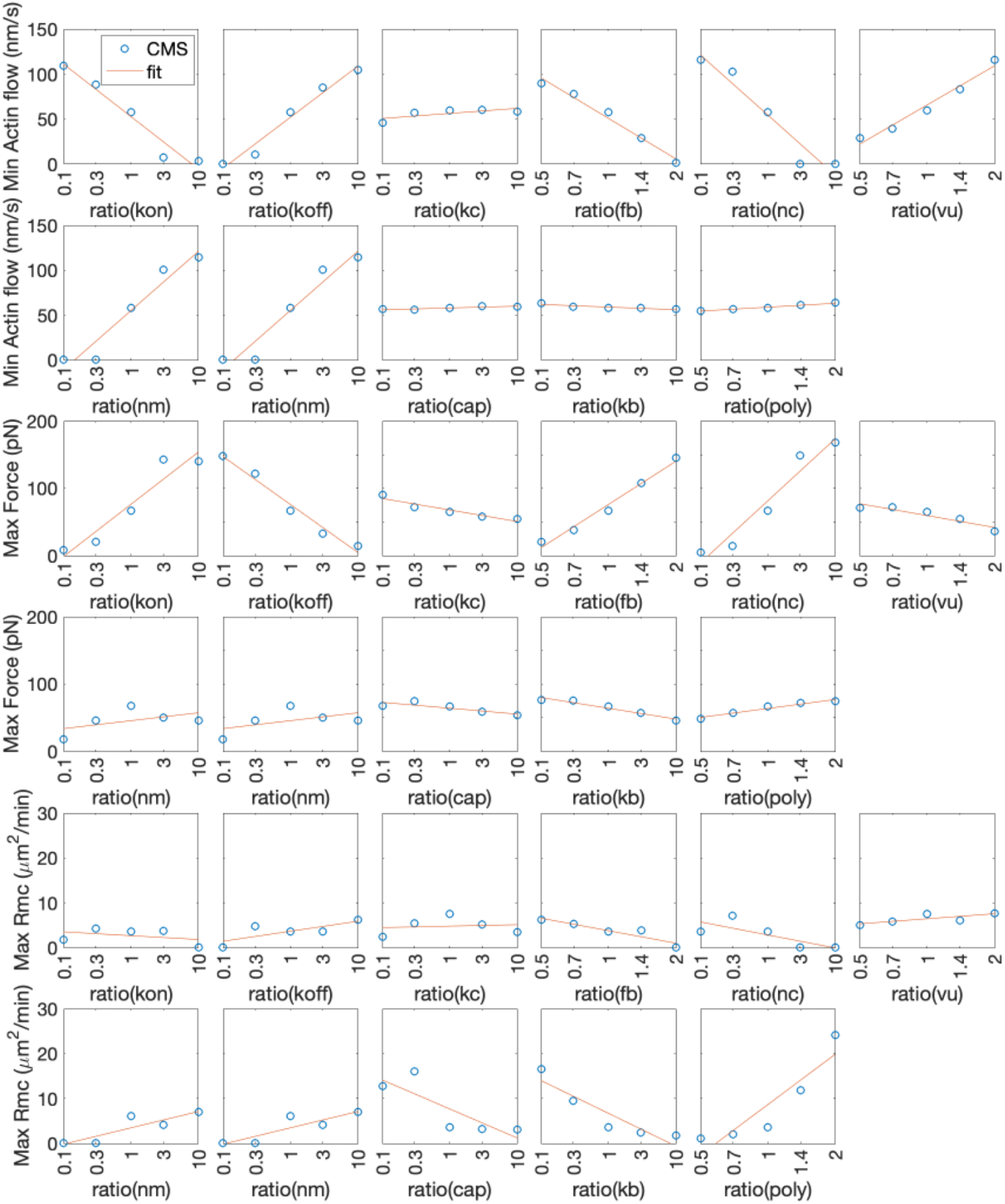
CMS parameter sensitivities. Predicted maximum RMC, maximum traction forces, and minimum F-actin flow over different stiffnesses were plotted with different parameter ratios compared to the base parameter values (Table 1). The linear regression between the migration predictions (Y axis) and the logarithm of parameter ratios (X axis) were plotted.

**Figure S2.**
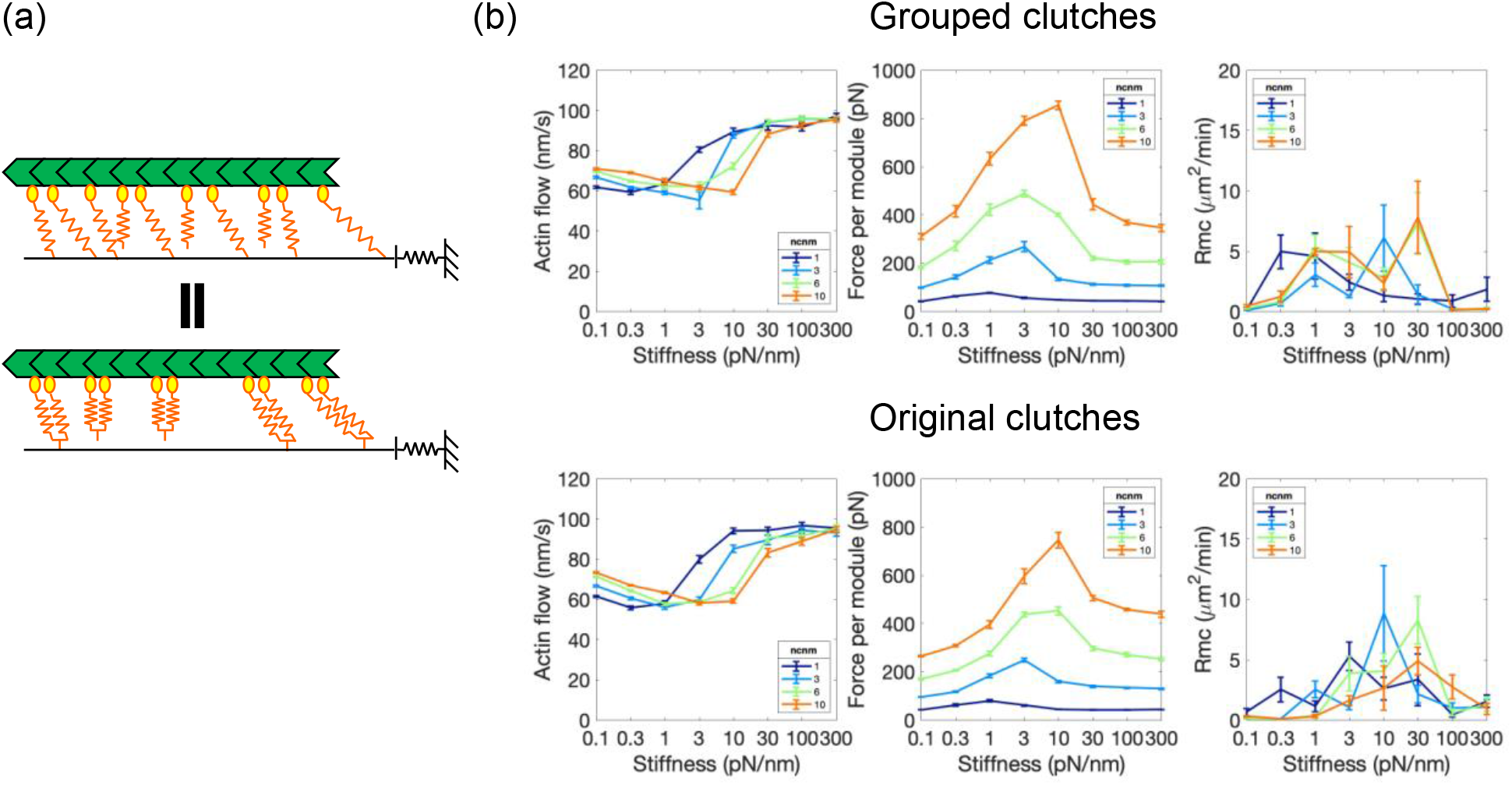
Grouped clutch approach enhanced the computational efficiency. (a) With sufficient clutch number, the system with grouped clutches (multiple clutches simulated as one clutch with stiffness multiplying the group clutch number, Equation 8) were equivalent to the system with individual clutches, thus enabling simulations larger clutch ensembles without increasing computational run time. (b) The CMS predictions with grouped clutches were equivalent to those without grouped clutches.

**Figure S3.**
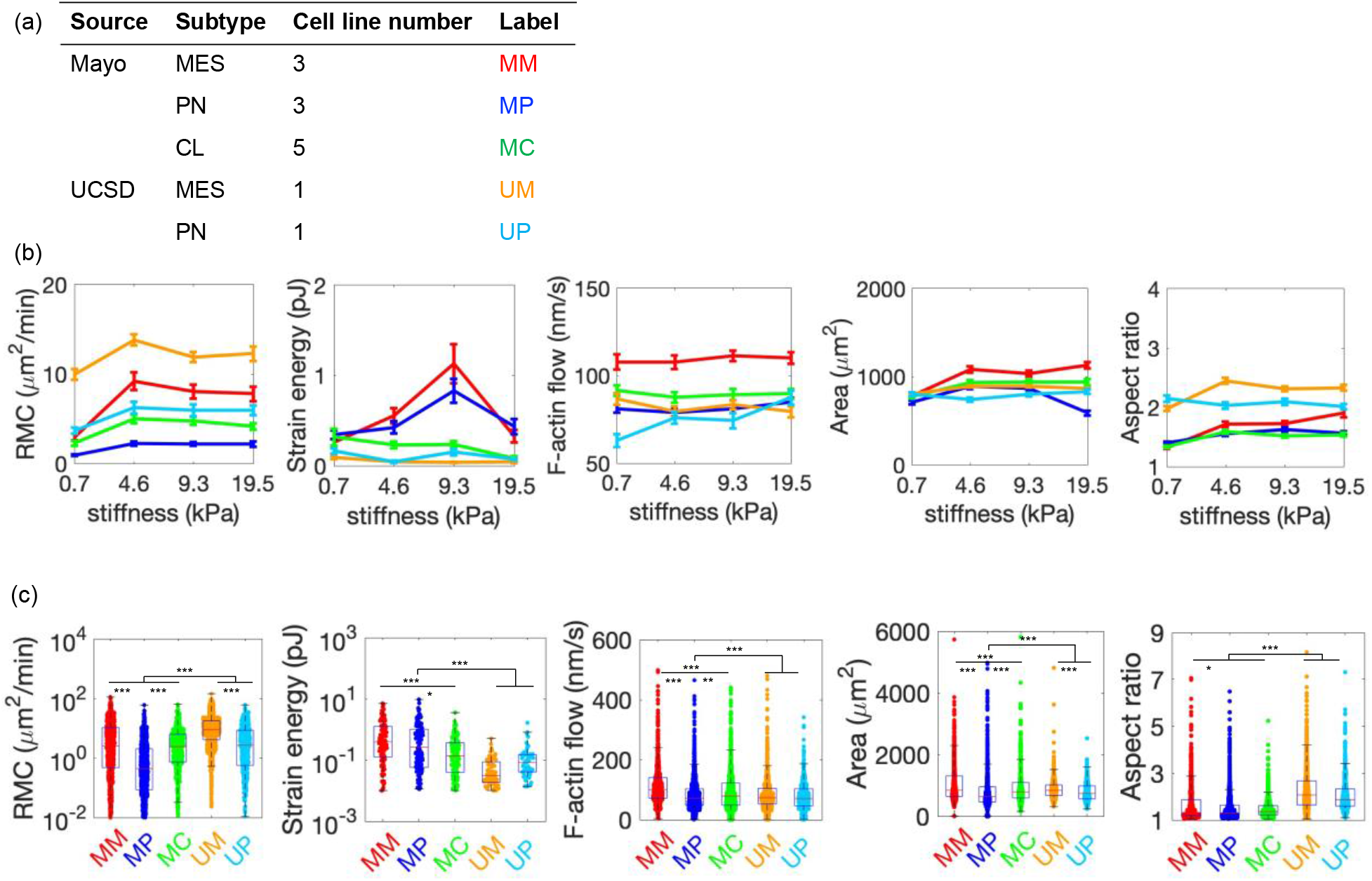
Heterogeneity in migration phenotypes of glioblastoma patient cells. (a) The experimental data set included 3 Mayo MES lines (MM), 3 Mayo PN lines (MP), 5 Mayo CL lines (MC), 1 UCSD MES line (UM), and 1 UCSD PN line (UP). (b) Mayo and UCSD cells were plated on polyacrylamide gels with stiffnesses of 0.7, 4.6, 9.3, 19.5 kPa, coated with laminin and Matrigel, respectively. Random motility coefficient (RMC), traction strain energy, F-actin retrograde flow, cell area, and aspect ratio were measured using established protocols [**Error! Reference source not found.**] and plotted for different substrate stiffnesses. (c) Data from all stiffness were aggregated. RMC, strain energy, and F-actin flow of Mayo MES, PN, CL cells and UCSD MES and PN cells were plotted with statistical significance indicated. * Denotes p<.05, ** p<.01, *** p<.001 derived from the Kruskal-Wallis test with Dunn-Sidak *post hoc* analysis.

**Figure S4.**
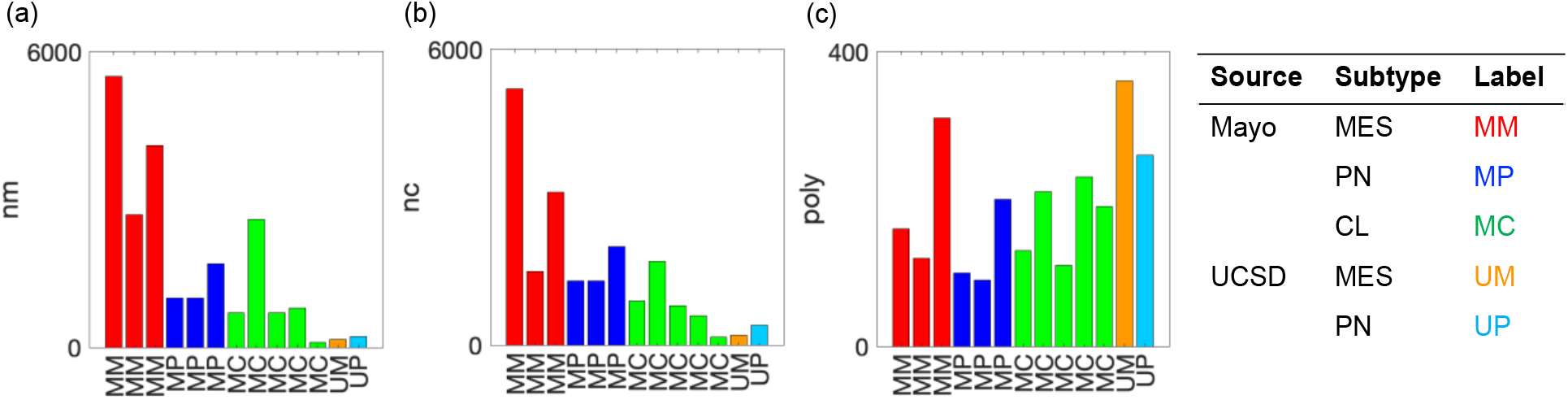
CMS-derived physical parameter values of glioblastoma patient cell migration. The motor number (a), clutch number (b), and F-actin polymerization rate (c) for each PD(X) line were plotted.

**Figure S5.**
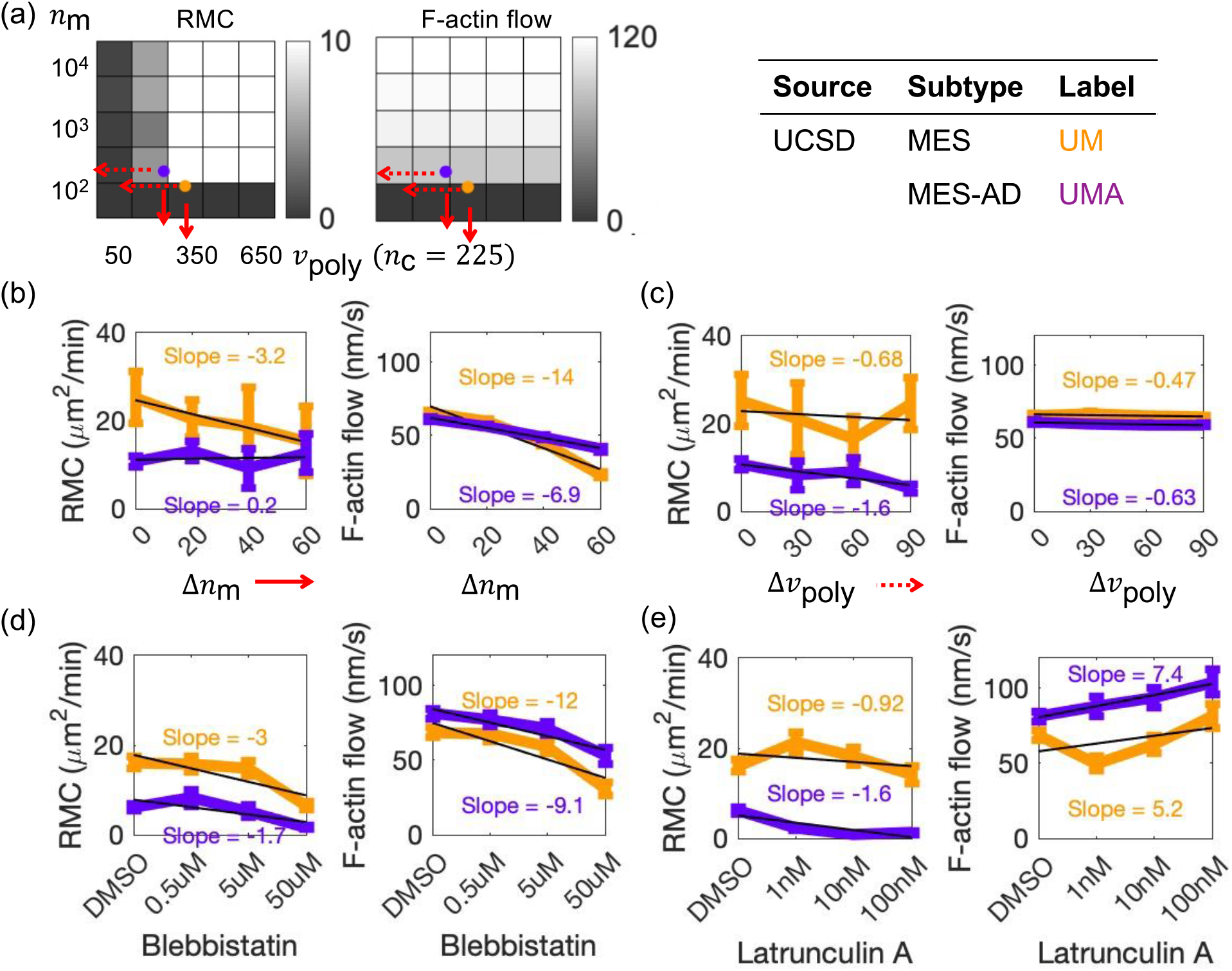
The CMS predicted UCSD MES and MES-AD cell migration and F-actin flow with actomyosin drug perturbations. (a) UCSD MES cells were cultured adherently for a week to become UCSD MES-AD cells before conducting migration assays. The heat map of the CMS prediction, including motility (RMC), traction force, and F-actin retrograde flow, with various values of (*n*_m_, *v*_poly_) with a constant value of *n*_c_ = 225. UCSD MES cells (orange) had higher F-actin polymerization rate and lower motor number compared to UCSD MES-AD cells (purple), which produced higher motility and higher sensitivity of F-actin flow with reducing the motor number. (b,c) The mean ± SEM values of RMC and F-actin flow of UCSD MES and MES-AD cells with decreasing motor number (b) (solid-red arrow) and decreasing F-actin polymerization rate (c) (dash-red arrow) predicted by the CMS were plotted. (d,e) The mean ± SEM values of RMC and F-actin flow of UCSD MES and PN cells with different concentrations of myosin II inhibitor, blebbistatin (d), and F-actin polymerization inhibitor, Latrunculin A (e), were plotted.

**Figure S6.**
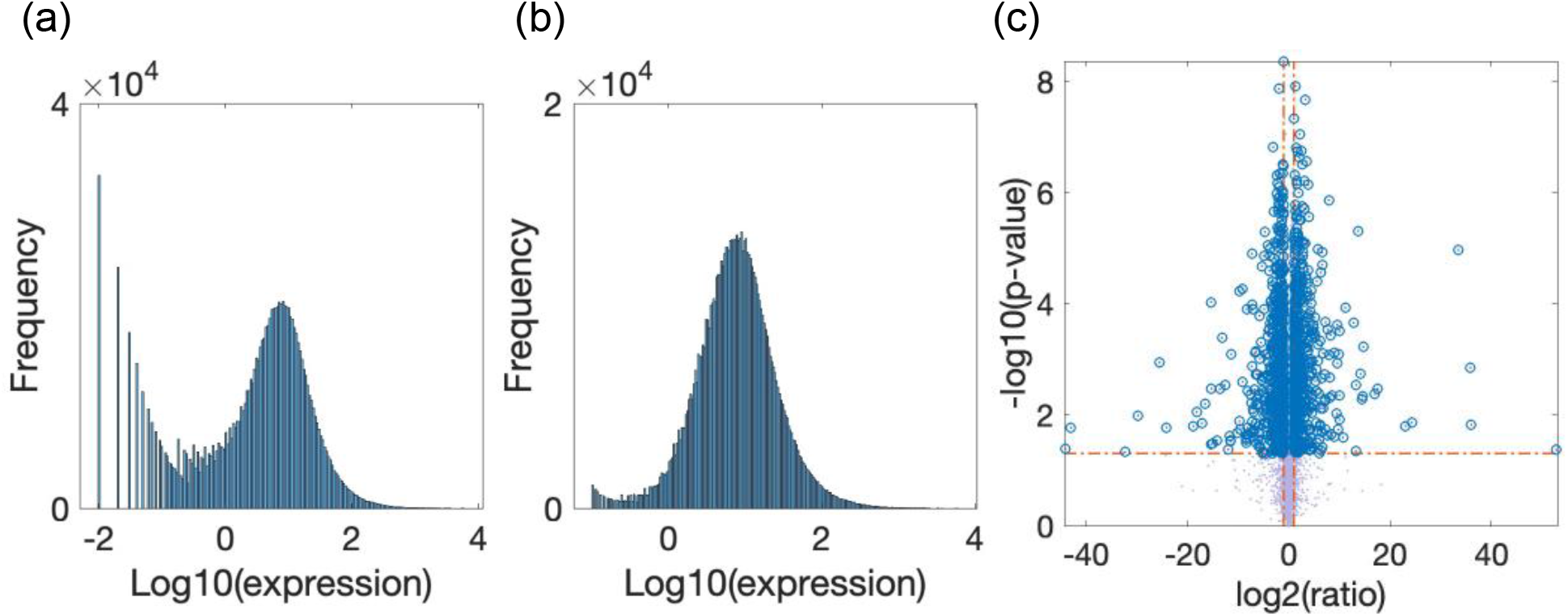
Differential migration genes between Mayo MES and PN lines. (a) RNAseq RPKM expression of the Mayo cells were used from Vaubel et al., 2020 [**Error! Reference source not found.**] (19,552 genes in 20 MES patients, 16 PN patients, 30 CL patients) with the non-Gaussian gene expression distribution. (b) Genes with geometric means less than 1 and with counts in less than 80% of patients were filtered out to achieve an approximately Gaussian gene expression distribution (11752 genes). A pseudo count of 0.1 was added to the expression level for further analysis to avoid small expression level (<0.1) (c) A two-sample t-test was applied to the mRNA expression of Mayo MES (20) and PN (16) lines, and the differential genes (1,175 genes) with p < 0.05 and fold change > 2 were obtained as shown in the volcano plot.

**Figure S7.**
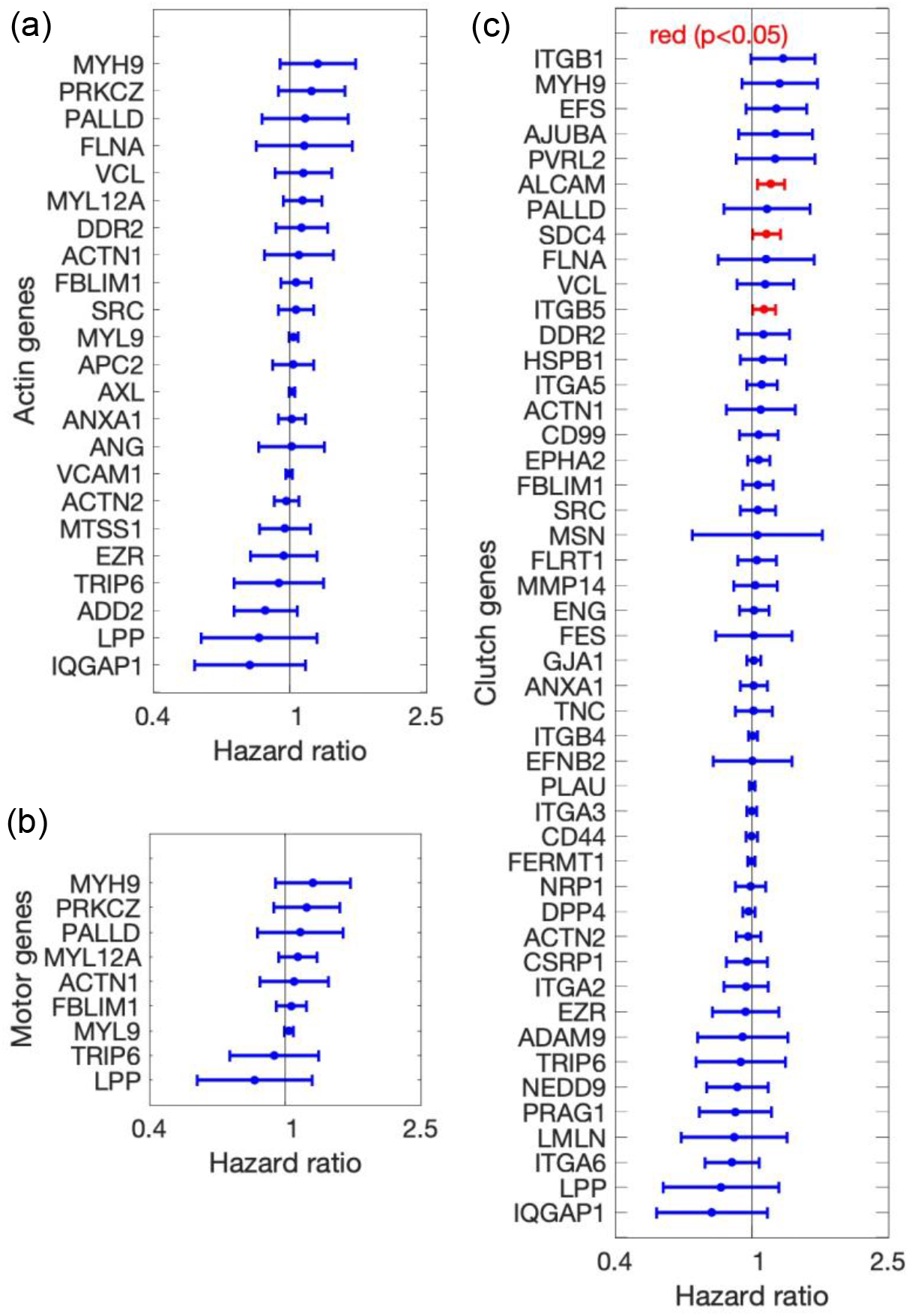
Cox regression analysis between gene expressions and patient overall survival (OS). We derived 23 actin genes (a), 9 motor genes (b), and 47 clutch genes (c) based on Gene Ontology cellular components from 1177 differential genes between Mayo MES and PN cells (Method: mRNA expression analysis). We applied the Cox regression analysis between the mRNA expression ratios of the actin (a), motor (b), clutch (c) genes in N=66 Mayo patients and their OS, and their hazard ratios with 95% Confidence Intervals were sorted and plotted in (a-c), with significant hazard ratios in red. The results show that the correlated genes with the OS is not different than that expected by chance, since the identified number of genes, 3, represents less than < 5% of 79 total genes).

